# The Dark Side of Photosymbiosis: Elaborate Repeats with TEs and Unknown ORFs in Photosymbiotic Bivalves Made Exceptionally Big Metazoan Mitogenomes

**DOI:** 10.1101/2025.01.13.632863

**Authors:** Andy D. Y. Tan, Ruiqi Li, Jingchun Li

## Abstract

Limited truly “complete*”* mitogenomes have been published for marine invertebrates, with many being partial and focused on protein-coding genes. Dispelling the traditional myth of “metazoan mitogenomes being highly conserved in size and gene order”, long-read technologies have revealed novel structures and complexities in animal mitogenomes. Our investigation of PacBio-assembled mitogenomes of several marine bivalve species (family Cardiidae), revealed a photosymbiotic bivalve, *Fragum sueziense*, possesses one of the largest circular metazoan mitogenome (92,770 bp). Additionally, mitogenomes of photosymbiotic bivalves from the genera *Fragum* and *Tridacna* ranged from 22 to 92.8 kb, surpassing the more typical sizes in non-photosymbiotic cardiids (∼14 to 19 kb). Those expansions in mitogenomes are attributed to elaborate, species-specific repetitive sequences in the major non-coding region (NCR) which manifest an inability to assemble “complete” mitogenomes from Illumina short reads (at 150 bp) alone; divergent nature of those NCRs also hinder interspecies alignment. However, we annotated supernumerary tRNAs, transposable element fragments and open reading frames in NCRs despite their hitherto unknown functions. We postulate that NCR inflation in these photosymbiotic species may be associated with elevated reactive oxygen species in their mantle and altered immune states due to host-symbiont interactions involving photosymbionts. More complete mitogenomes are needed to uncover novel genetic elements and their functions otherwise undocumented to science.

**SIGNIFICANCE STATEMENT:** Many published animal mitogenomes does not truly span the entirety of the major non-coding region (NCR), often due to the inability of short-read sequencing to confidently cover highly repetitive sequences. In this study, long-read sequencing enabled us to access “more complete” mitogenomes for a comparative analysis focused on the nature of the NCRs between photosymbiotic and non-photosymbiotic bivalves, and the elements within them. We not only uncovered one of the largest circular metazoan mitogenomes (*Fragum sueziense*), but also found ample genetic elements within the conventionally “non-coding” regions, including extensive elaborate repeat patterns, transposable elements and unknown open reading frames. To our knowledge, this is the first time extensive transposable elements have been reported in animal mitogenomes. Our study revealed extreme mitogenome expansions and complexities as potential costs to bivalve-algal photosymbiosis, and provide insights into metazoan symbiosis evolution.

## INTRODUCTION

Arisen as an endosymbiont of α–proteobacteria origin, mitochondria are essential organelles most known for their energy generation in the form ATP in eukaryotic cells (Roger *et al*. 2017). Owing to their abundance in animal tissues and relatively small size, mitochondrial DNA (mtDNA) and more recently mitochondrial genome (henceforth mitogenome), can be easily accessed by evolutionary biologists. Common practice in feature annotation and structural description of mitogenome since the advent of DNA sequencing in the 1980s has given rise to the ‘textbook description’ of the Metazoan mitogenome, based on vertebrates and model organisms: 37 invariantly arranged and compactedly ordered genes for 13 Protein Coding Genes (PCGs), 2 ribosomal subunit RNA genes (rRNAs) and 22 transfer RNA genes (tRNAs), with a non-coding ‘control region’, all nestled within 16k nucleotides that are strictly inherited matrilineally (Wolstenholme 1992; Boore 1999; Ghiselli, Gomes-dos-Santos, *et al*. 2021). Many studies involving phylogenetics (Plazzi *et al*. 2011; Liao *et al*. 2018), genetic diversity and populational variability (Gagnon *et al*. 2015; Hui *et al*. 2017) have long persisted with, and perpetuated such conventions established around mtDNA and/or mitogenomes. Mitochondrial gene rearrangement events are considered scarce and were used to infer deep evolutionary relationships at higher taxonomic levels (Boore 1999).

Outside the ‘textbook description’, many idiosyncratic exceptions, however, have been documented (Ghiselli, Gomes-dos-Santos, *et al*. 2021), including (1) non-clonal inheritance such as heteroplasmy (Kmiec *et al*. 2006; Wang *et al*. 2010), and doubly parental inheritance that thus far appear unique to certain Bivalvia lineages (Zouros 2013; Breton *et al*. 2021); (2) variations in number and location of genes, especially tRNAs (Boore 1999; Gissi *et al*. 2008); (3) variance and instability in structure and gene order outside of most deuterostomes (excepting tunicates; *17*); (4) variation in number, size and location of the so-called non-coding regions (NCRs) which include control regions (Larizza *et al*. 2002; Saccone *et al*. 2002; Wu *et al*. 2009).

Increasingly, researchers realize that these “exceptions” might be the norm, and our inability to find structural variations in the mitogenome is a result of technological limitations. Many of these novel structures and features in the Metazoan mitogenome can only be inferred from truly complete mitogenomes with confident characterization of major extensive NCRs (Xiaokaiti *et al*. 2022), which is enabled by long-read sequencing technologies (PacBio and Nanopore). Previously, these complex regions, particularly with extensive repeats, palindromes, and duplications have been challenging to assessed through target-enrichment or primer-walking Sanger sequencing (Kocher *et al*. 1989; Imanishi *et al*. 2013) and short-read whole-genome sequencing (Ekblom *et al*. 2014).

The Vertebrate Genomes Project Consortium’s reassessment of mitogenomes for 125 species using PacBio and Nanopore yielded “more complete” mitogenomes that are longer than those of the same species previously published as ‘complete’ reference mitogenomes on GenBank (Formenti *et al*. 2021). In light of novel gene region duplications and hypervariable repeats largely noted in the NCR(s) of vertebrate mitogenome (Formenti *et al*. 2021), we see the need for more extensive examinations of complete invertebrates mitogenomes. For instance, there are presently only ∼50 *complete* molluscan mitochondrial genomes sequenced through long-read technology that are available on GenBank (assessed May 2024), in contrast to the estimated 150,000 mollusc species in the ocean alone (Bouchet *et al*. 2016).

Some studies consider the control region the “longest NCR” in the Metazoan mitogenome (Larizza *et al*. 2002; Bronstein *et al*. 2018) while others report multiple origins of replications and regulatory elements in multiple control regions within NCRs (Jiang *et al*. 2007; Xiaokaiti *et al*. 2022). Generally, a control or displacement-loop (d-loop)-containing region, contain the origin of replication for either strand of the mitogenome, with the one for heavy strand replication often found in the major NCR of a mitogenome (Sbisà *et al*. 1997; Larizza *et al*. 2002). In this paper, we broadly define an NCR as a region of the mitogenome that does not encode the conventional 37 genes: PCGs, rRNAs and tRNAs; and the major NCR, being the longest NCR in each mitogenome. For us, an NCR, flanked by coding sequences, starts immediately after the last nucleotide encoding for a stop codon of the coding region, and ends a position before the start codon of another coding region. Biological functions of the mitochondrial NCR, especially in invertebrates, have not been thoroughly assessed. Studies in vertebrate systems have shown that the NCR contains certain conserved regions, representing sites for origin of mitochondria replication, transcription initiation, or other gene regulatory elements (Jemt *et al*. 2015; Sebastian *et al*. 2021). Some of these functions are achieved through the formation of secondary structures (Pereira *et al*. 2008; Jemt *et al*. 2015). Despite growing research on molecular architectures of metazoan NCR, our understanding of its sequence evolution and ecological role in animal adaptation remains minimal.

Given that mitochondria catalyze respiratory substrates oxidization, it is within reason to expect that animals living in extreme oxygen habitats (hypoxia or hyperoxia) exhibit unique mitochondrial gene regulations, as such environments may require dramatic changes in an animal’s cellular energy demands and metabolism (Hourdez and Lallier 2007; Welker *et al*. 2013). And since NCR may play an important role in mitochondria gene regulation, we may expect NCR sequence evolution to be significantly influenced by an animal’s environmental oxygen profile. In addition, extreme oxygen levels will assert high oxidative stress to the mitochondria (Xuefei *et al*. 2021) and increase DNA replication error and mutation rates, further impacting NCR evolution. A direct link between NCR evolution and extreme environmental oxygen level has not been established so far, but emerging studies assembling truly complete metazoan mitogenomes started to provide evidence supporting this hypothesis. For example, elapid sea snakes have low skin blood oxygen levels to support cutaneous respiration (Palci *et al*. 2019). Compared to their terrestrial relatives, sea snake mitogenomes contain much longer NCR regions characterized by tandem repeats and lack of conserved sequence blocks (Xiaokaiti *et al*. 2022). The deep sea chemosymbiotic bivalve *Calyptogena marissinica* also exhibits expanded tandem repeats and other special sequence elements in the NCR, which were hypothesized to aid in mitochondrial energy metabolism in the deep sea (Yang *et al*. 2019).

Photosymbiosis, where a heterotrophic organism hosts photosynthetic symbionts, presents a unique extreme oxygen microenvironment due to high levels of oxygen produced by symbiont photosynthesis. High rates of net photosynthesis create hyperoxia in the host during the day, while holobiont respiration results in hypoxia or even anoxia at night (Gardella and Edmunds 1999; Linsmayer *et al*. 2020). This extreme fluctuation in oxygen levels poses significant challenges to mitochondrial gene regulation. Therefore, it’s natural to ask whether NCR regions in photosymbiotic animals possess unique evolutionary signatures resulting from these challenges.

Thanks to recent large-scale genomic sequencing consortiums, such as the Aquatic Symbiosis Genomics (ASG) Project (McKenna *et al*. 2021), long-read mitochondrial genome assemblies of some photosymbiotic animals are now available. Compared to other photosymbiotic animals (*e.g.*, corals and sponges), bivalves cast less challenges in tissue preservation, DNA extraction, and library preparation for high quality genome assembly pipelines, allowing them to be the first several genomes being published through ASG (see Table 1 for genome note citations). With their ability to endure extreme oxygen fluctuation, and multiple complete mitogenomes available, photosymbiotic bivalves represent a great system to investigate the evolution of mitochondrial genomes and the NCR (Ghiselli, Iannello, *et al*. 2021).

**Table 1.** Information of all mitogenomes analyzed in this study. **PB:** PacBio Long-Read Sequencing; **IL:** Illumina Short-Read Sequencing; **PW-PCR:** Primer-Walking PCR; **ONT:** Oxford Nanopore Technologies. ***P***: photosymbiotic, ***NP***: non-photosymbiotic.

| GenBank Accession | Ecology | Species | Sequencing Technology | Genome Coverage | Citation |
| --- | --- | --- | --- | --- | --- |
| OX336384.1 | *P* | *Fragum fragum* (Linnaeus, 1758) | PB + Hi-C + IL | 14 | (Li, Li, Lemer, *et al.* 2024a) |
| OX411561.1 |  | *Fragum whitleyi* Iredale, 1929 | PB + Hi-C + IL | 43 | (Li, Li, Lemer, *et al.* 2024b) |
| OY804611.1 |  | *Fragum sueziense* (Issel, 1869) | PB + Hi-C | 34 | (Li *et al.* 2024) |
| OX031074.1 |  | *Tridacna crocea* Lamarck, 1819 | PB + Hi-C | 42 | (Li *et al.* 2023) |
| OY723432.1 |  | *Tridacna derasa* (Röding, 1798) | PB + Hi-C | 49 | (Li, Li, Lopez, *et al.* 2024a) |
| OX244045.2 |  | *Tridacna gigas* (Linnaeus, 1758) | PB + Hi-C | 36 | (Li, Li, Lopez, *et al.* 2024b) |
| NC_022194.1 | *NP* | *Fulvia mutica* (Reeve, 1844) | PW-PCR | - | (Imanishi *et al.* 2013) |
| NC_008452.1 |  | *Acanthocardia tuberculata* (Linnaeus, 1758) | IL | - | - |
| OX401423.1 |  | *Cerastoderma edule* (Linnaeus, 1758) | ONT+IL | - | - |

Extant, obligate photosymbiotic bivalves all belong to the family Cardiidae: giant clams (subfamily Tridacninae) and some heart cockles (subfamily Fraginae), mostly are shallow water dwellers (Li *et al*. 2020). The rest of Cardiidae includes non-photosymbiotic species, which are shallowly infaunal to epifaunal in tropical-subtropical areas (Herrera *et al*. 2015). The similar ecological niches of the photosymbiotic and non-photosymbiotic species make them ideal candidates for studying molecular evolution through comparative methods. For example, comparative transcriptomic studies have revealed unique molecular responses to reactive oxygen species (ROS) in a symbiotic bivalve (Li, Zarate, *et al*. 2024). Therefore, it would be particularly interesting to investigate how photosymbiosis influenced bivalve mitogenome evolution through a comparative lens.

This paper utilized photosymbiotic and non-photosymbiotic cardiid bivalves to reveal novel features and structures in metazoan mitogenomes. In particular, we aim to investigate whether unconventional NCRs are prominent in these bivalves, what are their potential functions, and whether the existence of novel NCRs are correlated with a photosymbiotic ecology.

## RESULTS

Complete mitogenomes of six photosymbiotic and three non-photosymbiotic, marine bivalve species from the family Cardiidae (Table 1) were verified, annotated and analyzed for general mitogenome parameters (genome and NCR sizes, gene annotations, order and rearrangements, repeat patterns, OCF and TE annotations). Phylogeny of focal species is depicted in Fig. 1, alongside representations of NCR:genome portion of the mitogenomes.

**Figure 1:**
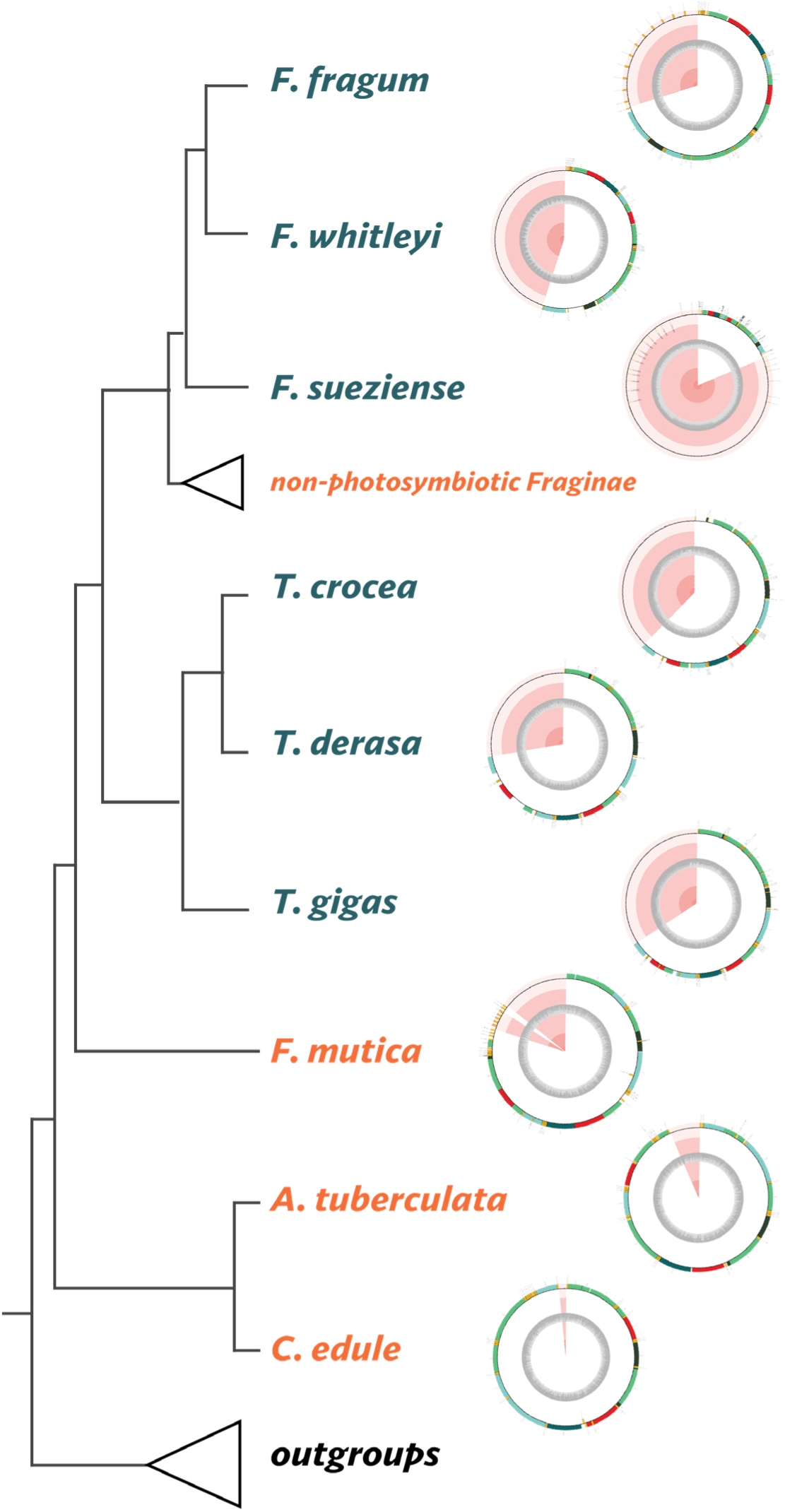
Circularized mitogenomes of cardiids shown with a cardiid phylogeny modified from Li et al. (2020). The region shaded red in the each mitogenome denote the position and proportion of the major NCR in relation to the whole mitogenome.

### Larger Genome Sizes and NCR Sizes in Photosymbiotic Species

The mitogenomes of cardiid bivalves are circular DNA molecules. Here we report *F. sueziense* having one of the largest known circular mitogenomes in metazoans, at 92,770 kbps. Instead of the typical 15-16 kbps expected for metazoan mitogenome sizes, the mitogenomes of the six photosymbiotic species span 22.4 to 92.8 kbps, with a median and mean of 27.1 kbps and 37.5 kbps respectively. A single major NCR > 6.6 kbps is found in all photosymbiotic species, and the average NCR size is 20.5 kbps (Table 2). Amongst the photosymbiotic species, the two largest major NCRs are found in *F. sueziense* (75.7 kbps) and *F. whitleyi* (13.8 kbps), making up to nearly 82% and 45% of the entire mitogenome respectively; whereas the smallest major NCR found is 6.6 kbps in *F. fragum*, which is 26.7% of its mitogenome.

**Table 2.** Comparison of mitogenome sizes, major NCR sizes, percentages of major NCR to full mitogenome sizes and repeat types found in each photosymbiotic and non-photosymbiotic cardiids in this study. ***P***: photosymbiotic, ***NP***: non-photosymbiotic.

| Species | Ecology | mtG size (bp) | Major NCR Size (bp) | NCR:Genome Percentage (%) | # of ORF (>75 nt) | Repeats |
| --- | --- | --- | --- | --- | --- | --- |
| *Fragum fragum* | *P* | 22,357 | 6,638 | 29.70 | 368 | c-SSRs-like |
| *Fragum whitleyi* | *P* | 30,336 | 13,752 | 45.33 | 580 | c-SSRs-like + satellite block |
| *Fragum sueziense* | *P* | 92,770 | 75,654 | 81.55 | 924 | undescribed |
| *Tridacna crocea* | *P* | 29,009 | 11,083 | 38.21 | 476 | undescribed |
| *Tridacna derasa* | *P* | 24,955 | 7,110 | 28.49 | 402 | undescribed |
| *Tridacna gigas* | *P* | 25,335 | 8,785 | 34.68 | 381 | undescribed |
| *Fulvia mutica* | *NP* | 19,110 | 2,612 | 13.67 | 293 | 3 minisatellites |
| *Acanthocardia tuberculata* | *NP* | 16,104 | 1,101 | 6.84 | 281 | 1 minisatellite |
| *Cerastoderma edule* | *NP* | 14,943 | 267 | 1.788 | 231 | Absent |

The non-photosymbiotic species used for comparison in this study have mitogenomes that range from 14.9 to 19.1 kbps. In contrast to photosymbiotic species, major NCRs, if present in non-photosymbiotic cardiids, occupy only up to 13.7% of their mitogenomes. The largest major NCR identified for non-photosymbiotic species is 2.6 kbps in *F. mutica*. The major NCR in *C. edule* has only 267 bp. For more comparisons of descriptive parameters between photosymbiotic and non-photosymbiotic mitogenomes, see Fig. S1.

### Additional Verification of NCR Presence/Absence

PacBio long read sequencing revealed major NCR with extensive long repeats (>200 bp) in the mitogenomes of all photosymbiotic bivalves sequenced. Reference-guided assembly mapping Illumina short reads (150 bp) to PacBio mitogenome for *F. fragum* and *F. whitleyi* confirm the presence and length of the major NCR based on read and repeat coverage (>350 bp over >11 times in both species). For both species, we observe higher than mean coverage for their major NCR region, which contains repetitive motifs that challenges mapping, leading to ambiguous assembly and failure to fully retrieve true identity of said repeated region.

Meanwhile, short or absence of major NCR(s) in non-photosymbiotic cardiids is unlikely a sequencing or assembly error. Mitogenomes for the non-photosymbiotic species in our study are sequenced through three distinct sequencing techniques: (1) coupling Nanopore to Illumina for *C. edule* presents long sequences that would cover the entirety of the NCR, hence bridging the coding regions; (2) careful primer-walking PCR of 21 *F. mutica* individuals manage to span through the entire, albeit short, NCRs (Imanishi *et al*. 2013); (3) while the mitogenome of *A. tuberculata* was sequenced with only Illumina short reads, their NCR yield similar raw read coverage to the core coding region, not with higher coverage and uncertain mapping; Given these outcomes from the three techniques, short NCRs in non-photosymbiotic cardiids is highly probable.

### Mitogenome Composition

By gene annotation, we identified for each species the complete set of standard 37 mitochondrial genes —13 PCGs (*cox1*-3, *atp6*, *atp8*, *nad1-6*, *nad4l, cytb*), 2 rRNAs (*lrRNA* and *srRNA*) and 22 tRNAs (including two copies of *trnL* and *trnS*; Table 3). *atp8*, which has been repeatedly reported absent in bivalve mitogenomes (Serb and Lydeard 2003; Liao *et al*. 2018; McElroy *et al*. 2024), is annotated for all species in this study. Consistent with recent analyses finding *atp8* in various bivalves (Plazzi *et al*. 2016), including but not limited to Mytilidae (Plazzi *et al*. 2016; Zhao *et al*. 2022), Pectinidae (Malkócs *et al*. 2022), our finding supports the presence of *atp8* sequences in Cardiidae, and flags the need for researchers to attempt deeper search for *atp8* gene if applicable (Gan *et al*. 2019), and exercise caution when making further proclamation that a mitochondrial gene is lost.

**Table 3.** Summary table of genes annotated for all species.

| Species | PCG | rRNA | OH | OL | miRNA | sRNA | snoRNA | tRNA | additional tRNA copies |  |  |  |  |
| --- | --- | --- | --- | --- | --- | --- | --- | --- | --- | --- | --- | --- | --- |
|  |  |  |  |  |  |  |  |  | M | N | S | C | other (x1) |
| *Fragum fragum* | 13 | 2 | - | - | 23 | 15 | 1 | 35 | 1 | - | 11 | - | H |
| *Fragum whitleyi* | 13 | 2 | - | 21 | 136 | 19 | 3 | 26 | 1 | - | 1 | - | L, Y |
| *Fragum sueziense* | 13 | 2 | 1 | 31 | 131 | 32 | 1 | 46 | 1 | 10 | 11 | - | G, I |
| *Tridacna crocea* | 13 | 2 | - | - | 28 | 10 | 3 | 25 | 1 | - | - | - | F, L |
| *Tridacna derasa* | 13 | 2 | 15 | - | 61 | 29 | 30 | 23 | 1 | - | - | - | - |
| *Tridacna gigas* | 13 | 2 | - | - | 211 | 16 | 21 | 27 | 1 | - | - | - | I, L, N, R |
| *Fulvia mutica* | 13 | 2 | 1 | - | 123 | 7 | 12 | 30 | 2 | - | - | 4 | F, I |
| *Acanthocardia tuberculata* | 13 | 2 | - | 1 | 6 | 6 | 3 | 23 | 1 | - | - | - | - |
| *Cerastoderma edule* | 13 | 2 | 1 | - | 13 | 4 | 3 | 26 | 1 | - | - | - | A,G,T |

With regard to tRNA genes, every cardiid species in this study has an additional copy of *trnM*, except for *F. mutica* which has 2 additional copies of *trnM*. We annotated three to 24 more additional copies of tRNAs for all species, except for *T. derasa* and *A. tuberculata*. We report two photosymbiotic *Fragum* spp. showing exceptionally high tRNA duplications in their major NCR. In *F. fragum*, 11 supernumerary copies of *trnS* were annotated in an evenly distributed manner along the c-SSRs-like repeats within its major NCR, on top of the standard two copies in its coding region (Fig. 4A). In *F. sueziense*, 10 additional copies *trnN* and 11 additional copies of *trnS* are annotated in the major NCR. These tRNA duplications events have not been previously reported in the literature. 5 copies of *trnC* are also detected in the NCR of non-photosymbiotic *F. mutica*, consistent with Imanishi *et al*. (2013)(Imanishi *et al*. 2013).

### High Levels of Mitochondrial Gene Rearrangements within Cardiida

A large number of gene rearrangements have been reported in bivalves (Hoffmann *et al*. 1992; Boore 1999; Malkócs *et al*. 2022), and our study supports the phenomenon not only for minor genome rearrangements (tRNAs only) but also major (PCGs and rRNAs) rearrangements (Fig. 2). Intrageneric gene order is slightly more conserved than that between genera. Between Fraginae and Tridacninae, we observe three positions for *cox2*, and only two positions for *nad3, nad4, nad4l* and *atp8*. Between photosymbiotic and non-photosymbiotic cardiids, however, gene arrangements appear not conserved and would require further synteny analyses, which is beyond the scope of this study.

**Figure 2:**
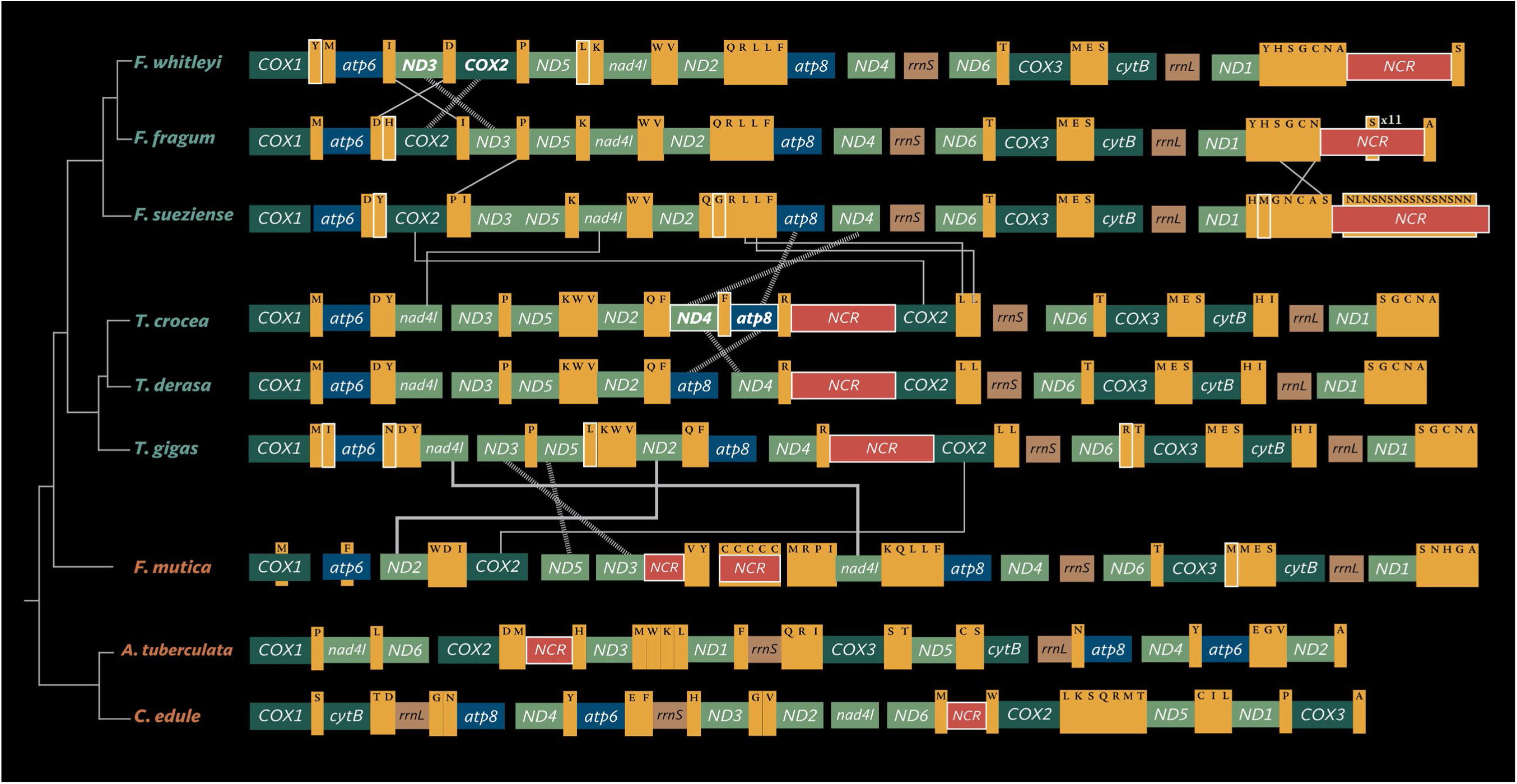
Mitochondrial gene rearrangement comparisons. PCGs in light green (nad genes), teal (cox genes) and blue (atp genes). Red for NCR, brown rRNAs and yellow tRNAs..

The placements of the major NCRs are consistent within smaller genera in Cardiidae but are inconsistent between genera. We found five loci where the major NCRs arise within cardiidae. Within Fraginae, the major NCR is found between *nad1* and *cox1* while it’s between *atp8/nad4* and *cox2* in Tridacninae. In *F. mutica*, the major NCR is between *nad3* and *nad4l.* The five placements suggest five possible origination events or lesser NCR rearrangement events likely prior to the expansion of NCRs within the photosymbiotic Fraginae and Tridacninae, similar to reports of multiple independent origins of inflated NCR with repetitive elements in the mitogenomes of various Pectinid lineages (Gjetvaj *et al*. 1992).

### Repeat patterns in the major NCRs

The major NCRs in these bivalve species mostly consist of sequence repeats. The types of repeat patterns found within the major NCRs are listed in Table 1 (also visualized in Fig. 4). Repeats in non-photosymbiotic species, if present, are minisatellites repeat pattern, and are simpler compared to those found in photosymbiotic species (Fig. 3).

**Figure 3:**
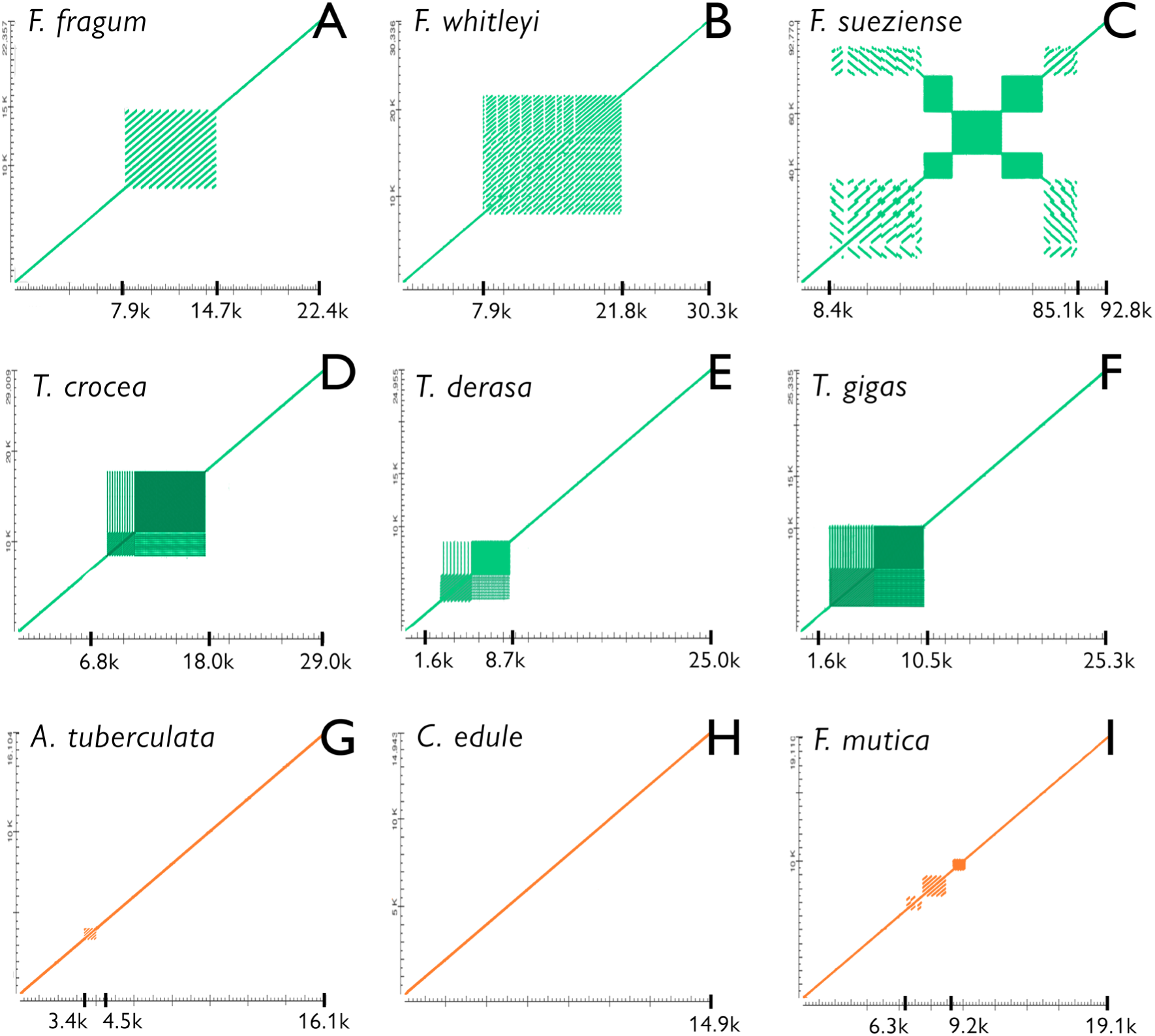
Dot plots for each mitogenome. Representative dot plots showing complexities of recurrent patterns in repeated sequences observed in the major NCRs of photosymbiotic (**A-F**) and non-photosymbiotic (**G-I**) cardiids.

**Figure 4:**
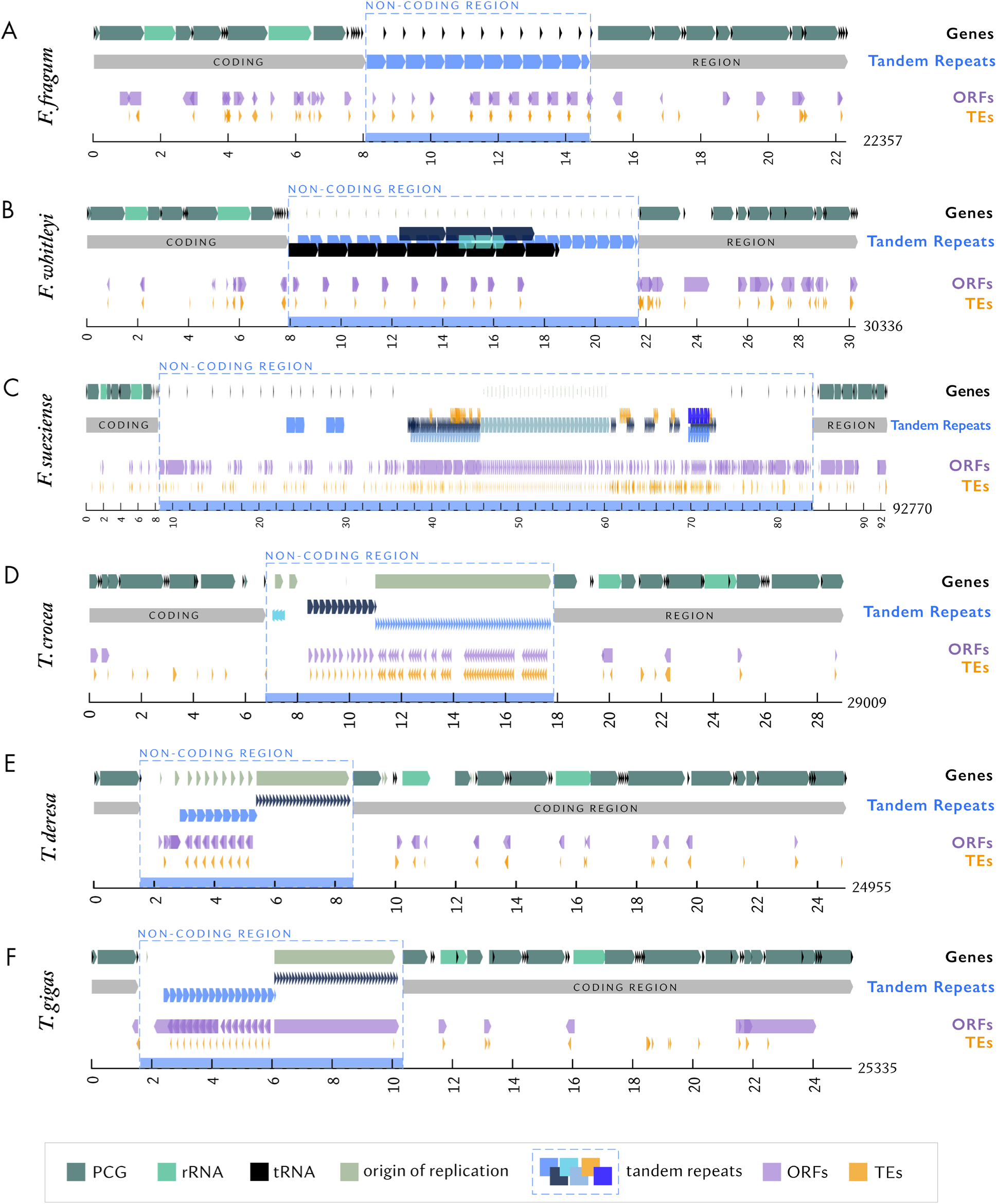
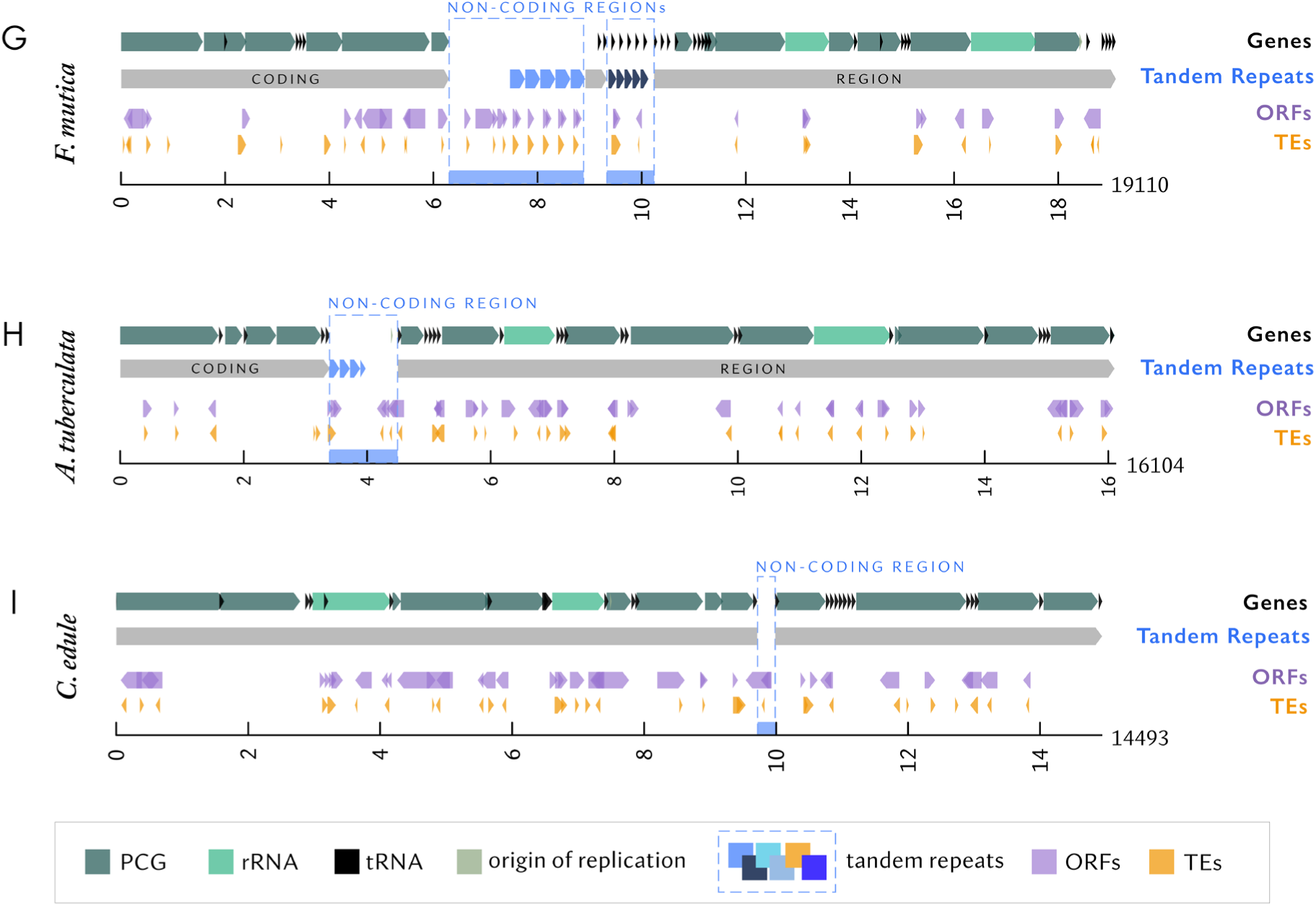
Linearized visualization of gene annotations for the mitogenomes. For each species, the first row visualizes PCGs and origin of replications, second row denotes pronounced tandem repeat patterns, third and fourth rows for ORFs and TEs that overlap with one another respectively. **A-F**: Photosymbiotic cardiids. For each species, the first row visualizes PCGs and origin of replications, second row denotes pronounced tandem repeat patterns, third and fourth rows for ORFs and TEs that overlap with one another respectively. **G-I**: Non-photosymbiotic cardiids.

The repeat pattern in the major NCR of *F. fragum* resembles a compound short sequence repeats (c-SSRs) described in plant mitogenome (Xiong *et al*. 2022). In a c-SSRs in plant, the repeat pattern typically has a “(repeats motif)*_n_* — species specific sequence — (repeats motif)*_n_*” form. In the case of *F. fragum*, we observe one ∼350 bps sequence motif repeated over 11 times, but each repeat motif is separated by >100 bps of individual specific random sequences [ in the form of “(repeat motif— individual/species specific sequence)*_n_*” ] (Fig. 3D). We are tentatively calling this c-SSRs-like repeat patterns. These c-SSRs-like repeats appear to have small proportions of interspecies transferability, and are expected to be overlooked between two closely related species (Xiong *et al*. 2022).

*F. whitleyi*, *F. sueziense* and *Tridacna* spp. have more complex repeat patterns that appear to have elements of a c-SSRs-like repeat and satellite blocks (Fig. 3B-F; see also Fig. 4 for visualization of selected tandem repeat patterns). Longer repeat motifs within their c-SSR-like sequences contain various smaller tandem repeats of various period length and copy numbers (see Data S2). Repeat types are more conserved within *Tridacna* than within *Fragum.* To our knowledge, these types of repeat patterns have never before been described.

### Transposable Elements in the mitogenomes and ORFs in major NCRs

We detected TE fragments that encode between 9 and 109 codons (median: 25 codons; mode: 22 codons) using Interspersed Repeat Protein Masking Based on Protein Similarity on RepeatMasker (Data S2). The TE sequences found are incomplete fragments, with many found to coincide with ORFs with between 78 and 4158 nucleotides (Fig. 4). Non-photosymbiotic cardiids each have 35 total TE fragments masked in their mitogenomes but the overall count within the NCR is less than 11 respectively (Table 4); photosymbiotic species have slightly elevated numbers of total TE (43 to 47). *F. sueziense* have very high count of TE fragments at 355 (318 in major NCR), followed by *T. crocea* at 94 (68 in major NCR).

**Table 4.** Summary table of TE fragment counts by types. The first six columns after species are most prevalent types detected.

| Species | LTR/<br>Copia | LTR/<br>Gypsy | LTR/<br>ERV-type | LTR/<br>Pao | LINE/<br>L1 | RC/<br>Helitron | LTR | LINE | DNA<br>transposon | TE in<br>NCR | Total<br>TE |
| --- | --- | --- | --- | --- | --- | --- | --- | --- | --- | --- | --- |
| *Fragum fragum* | **16** | **16** | 0 | 1 | 4 | 1 | 35 | 7 | 4 | 18 | 47 |
| *Fragum whitleyi* | 8 | **15** | 4 | 2 | 4 | 1 | 29 | 10 | 3 | 9 | 43 |
| *Fragum sueziense* | **79** | **61** | 7 | 25 | 16 | 9 | 195 | 56 | 9 | 318 | 355 |
| *Tridacna crocea* | **19** | 5 | 2 | 3 | **56** | 0 | 29 | 57 | 8 | 68 | 94 |
| *Tridacna derasa* | 6 | 5 | 1 | 1 | 1 | **9** | 13 | 4 | 4 | 12 | 30 |
| *Tridacna gigas* | 6 | 6 | **17** | 2 | 1 | 0 | 32 | 5 | 7 | 20 | 44 |
| *Fulvia mutica* | 7 | 5 | 3 | 3 | 2 | 0 | 35 | 12 | 3 | 10 | 35 |
| *Acanthocardia tuberculata* | 10 | 4 | 1 | 5 | 3 | 1 | 20 | 9 | 5 | 4 | 35 |
| *Cerastoderma edule* | 9 | 6 | 2 | 2 | 2 | 1 | 21 | 8 | 5 | 1 | 35 |

Long Terminal Repeats (LTR) retrotransposons or Long Interspersed Nuclear Elements (LINEs) are the most masked TE fragments integrated in these cardiid mitogenomes, with LTR/*Copia*-type, LTR/*Gypsy*-type and LINE/*L1*-type being the three most rampantly inserted TEs (Table 4). Unlike in *Arabidopsis* where *Ty1/copia*, *Ty3/gypsy* and *LINE*-like retrotransposons are scattered through the mitogenome without “preference for hotspots of integration” (Knoop, 1996), we find high copies of one or two types of TE fragments in consistently repeated chunks in the extended major NCR of photosymbiotic cardiid mitogenomes, making said retrotransposons the only or dominant type in those major NCRs. LTR/*Gypsy*-type TE is dominant in the NCR of all *Fragum* species. LTR/*Copia*-type TE, despite being the most commonly detected TE type in our analysis, is co-dominant with LTR/*Gypsy*-type TE for only *F. fragum* and *F. sueziense*, and with LINE/*L1*-type TE for *T. crocea*. The single dominant TEs in *T. derasa*, *T. gigas* and *F. mutica* are RC/*Helitron*-type, LTR-*ERVK-*type and LINE/*Ambal*-type TEs respectively. Without highly repeated major NCR, *A. tuberculata* and *C. edule* have no highly concentrated copies of a single dominant TE type. *Gypsy*-like and *Copia*-like LTRs are commonly detected in eukaryotic nuclear genomes and most abundant in plant genomes (Natali *et al*. 2015); it is hence interesting to find them in repeated copies, despite fragmentary, in metazoan mitogenomes. Given our study is only restricted to matching sequences to an existing TE database through RepeatMasker, more precise and comprehensive identification through *de novo* annotation might yield a more accurate count and identity estimates (Rodriguez and Arkhipova 2023).

All dominant TE sequences found in the NCRs appear fragmentary based on sequence length matches, likely either from formerly complete insertions or originated from partial duplications (Knoop *et al*. 1996). While it is unclear whether each repeated TE fragment within each NCR region degenerated concomitantly, or the duplications happened after a single degeneration event, the latter progression is more parsimonious.

## DISCUSSION

### Repetitive Elements and Gene Duplications in Expanded NCR as the Norm

Long-read sequencing can retrieve traditionally “non-coding regions” that were previously undetected, or unbridgeable due to long repetitive elements, thereby leading to assemblies of “more complete” reference mitogenomes. Comparing, for example, our *T. crocea, T. derasa* and *T. gigas* mitogenomes spanning ∼29 kbp, ∼25 kbp, ∼25 kbp respectively with >7k-long NCRs each, to purportedly complete reference genomes from short-read sequencing for mitochondrial genomes of the same species at 18,266 bp, 20,760 bp and 19,558 bp respectively (Ma *et al*. 2018; Ma *et al*. 2020; Baeza *et al*. 2022), the single inflated major NCR is the primary contributor to the larger sizes of photosymbiotic cardiid mitogenomes. Recent findings from many mitogenomes of vertebrate systems concur with observations of previously unreported repetitive elements and gene duplications in the “non-coding regions” similarly contributing to larger-than-conventionally- expected mitochondrial genome sizes (Formenti *et al*. 2021), though the extent of NCR expansion in photosymbiotic cardiids is even greater. Having an expanded NCR may not be the exception but norm in metazoans (Wang and Lavrov 2008; Formaggioni *et al*. 2021). We, therefore, recommend reconsiderations of completeness in presently published references, particularly through those from short-read sequencing technologies, and advocate for the use of long-read sequencing, when possible, for truly complete mitogenome sequencing.

### NCR and Its Implications in Bivalve Phylogenetics

Photosymbiotic bivalves from two separate cardiid subfamilies exhibit exceptionally long NCRs compared to non-photosymbiotic cardiid species. While Fraginae and Tridacninae are sister clades, the evolution of photosymbioses in both linages are considered independent (Li *et al*. 2020). The emergence of an extensive major NCR could be a plesiomorphic to for both subfamilies and *F. mutica*. Although NCRs are structurally similar within the genus *Tridacna* with significant overlap in repeat patterns, those identified in *Fragum* appear species-specific, likely fast evolving within the genus. Despite finding high gene rearrangements in cardiids, which seem prevalent in bivalves (Plazzi *et al*. 2016), conserved gene order within genera coupled with the lack of interspecific TE fragment similarities and fast turnover of species-specific NCR sequences might be useful in phylogenetic analyses for species delimitation.

### Excluding Doubly Uniparental Inheritance (DUI) as Potential Smokescreen

Some bivalve species exhibit DUI, a unique mitogenome inheritance mechanism, which may impact NCR evolution. In species with DUI, maternally transmitted (female-type) mitochondria exist in both genders, while paternally transmitted mitochondria (male-type) are predominantly in males, resulting in male heteroplasmy (Ghiselli, Gomes-dos-Santos, *et al*. 2021). Extensive work has been done on the mitogenomes of these species, revealing that they have large NCRs with complex genetic elements such as repeats and secondary structures, and the NCRs have gender specific functions (Zouros 2013; Guerra *et al*. 2014). Some of the NCR structures are a result of duplication and recombination events between the different types of mitogenomes in heteroplasmic males (Cao *et al*. 2009). It has also been hypothesized that the male-type mitogenome is experiencing relaxed selection, and thus evolve faster (Stewart *et al*. 1996). Despite the impacts of DUI in NCR evolution, it is unlikely to be the driver of the complex NCR structures in photosymbiotic cardiids, as DUI has never been reported in any species belonging to the family Cardiidae (or the order Cardiida). We also did not observe any evidence of strong heteroplasmy in the miteogenome assemblies, supporting that DUI was not present in species studied here.

Excluding the life history implications of DUI, rapid evolution of NCRs in photosymbiotic cardiids (and lack thereof in non-photosymbiotic species) is likely influenced by their unique symbiotic lifestyle. Here we posit three non-mutually exclusive hypotheses that could potentially explain the expansion of NCRs in the mitogenomes of photosymbiotic cardiids:

### Hypothesis I: Latent Effects of ROS and UV in Inducing Replication Errors

One of the most relevant ecological factors here is the higher-than-usual oxidative stress caused by the presence of algal photosymbionts. Mitogenomes are already sensitive to oxidative stress due to the lack of protective histones and the proximity to reactive oxygen (ROS) production sites (Starkov 2008). Such is amplified in photosymbiotic bivalves, because their tissues are constantly exposed to high oxygen levels resulting from algal photosynthesis. Their symbiont-hosting tissues also endure prolonged sun exposure to sustain ample light supply to the algae. Therefore, the hosts’ mitochondria may experience extensive UV and ROS-induced damages such as ATP depletion (He *et al*. 2022). Although photosymbiotic hosts utilize certain mechanisms (such as shell/tissue optical arrangement; *79*, *80*) to reduce UV and ROS impact, it is likely that these stresses still induce higher DNA replication errors and interfere with DNA repair in the mitochondria, which could be responsible for the observed highly variable and species-specific repeat sequences.

### Hypothesis II: Suppressed Immune System Lacking in Ability to Curb TE Proliferation

Rapid expansion of repetitive elements is not limited to the mitochondrial genomes of photosymbiotic cardiids; similar patterns have also been observed their nuclear genomes (Li 2024), indicating that oxidative stress alone is likely not the sole factor for the rapid expansions of NCR. Expansion of repetitive element have been identified in the nuclear genomes of several other photosymbiotic animals. For example, all three species of giant clams examined (*T. maxima, T. gigas, T. crocea*) exhibit approximately 70% repeat content, one of the highest known among bivalve (Li 2024). Within these species, TEs constitute the majority of the repetitive elements. This pattern is not restricted to bivalves alone. In zoantharians, for instance, the non-symbiotic species *Palythoa mizigama* and *Parazoanthus swiftii* exhibit a lower proportion of TEs compared to their symbiotic counterparts (Fourreau *et al*. 2023). This pattern is also observed in anemones and stony corals (Baumgarten *et al*. 2015). What might be unique about symbiotic organisms that facilitates the expansion of repetitive elements?

One possible explanation is their adapted immune systems, which have evolved to accommodate symbiotic lifestyles. Down-regulation of immune-related functions during the establishment and maintenance of symbiosis has been observed in various animal hosts, including corals (Pinzón *et al*. 2015) and heart cockles (Li, Zarate, *et al*. 2024). The immune system is known to play a role in TE suppression. For example, epigenetic modifications such as DNA methylation are involved not only in immune responses to bacterial infections (Qin *et al*. 2021) but also in TE silencing (Hollister and Gaut 2009). During and after the establishment of symbiosis, it is possible that the host’s immune system is suppressed long-term to sustain a stable relationship. Furthermore, prevalent symbionts like Symbiodiniaceae are also known to modulate the host’s immune system (Bove *et al*. 2022). Therefore, the suppression of host immune system could simultaneously impair TE silencing mechanisms, leading to the proliferation of TEs in both nuclear and mitochondrial genomes.

### Hypothesis III: NCR Exhibits Functional Value and Was Selected for

Substantial sizes of truly complete mitogenomes contradict the conventional notion that metazoan mitochondria are compact (*2*, but also see *68*) –– shorter genomes are advantaged with shorter time and lower higher energy requirement for their replication (Diaz *et al*. 2002). From an evolutionary perspective, natural selection would minimize energy costs by reducing the mitogenome size. However, this is not observed in photosymbiotic cardiids. This raises an intriguing question: why hasn’t natural selection acted to reduce the size of these mitogenomes despite the increased energy demands?

Replication error and TE insertion can introduce sequence repeats into the hosts’ mitogenomes, which are often eliminated or rendered non-functional through purifying selection. However, this our observations suggest long-term maintenance of these sequences. It is possible that the error and insertion rates are so high that the organisms cannot effectively suppress them, and the benefit of photosymbiotic association surpass the energy burden of replication large mitogenomes, so this tradeoff is tolerated. However, another distinct possibility is that such repeats and insertions exhibit functional value for the hosts, especially at the regulatory level.

Our annotation of the NCRs identified various additional RNA components in the mitogenomes of some photosymbiotic cardiids. For example, supernumerary tRNAs are positioned within the NRCs of *F. fragum* and *F. sueziense.* These are not false positives, because the sequences possess the canonical cloverleaf secondary structures (Laslett and Canbäck 2008). Although mitochondrial tRNA duplication, rearrangement, and gene model changes are not rare in Metazoa (Cantatore *et al*. 1987; Rawlings *et al*. 2003), possessing 11 additional copies of the same tRNA sequence (*F. fragum*) is still highly unusual. tRNAs are crucial for mitochondrial gene translation and many organisms need to import nucleus-encoded tRNAs to facilitate the process (Salinas-Giegé *et al*. 2015). Additional tRNA repeats in the Photosymbiotic bivalves could confer more efficient mitochondrial gene translation and may even accelerate mitochondria replication. Such mitochondria could possess adaptive advantages in the hyperoxia tissue environment, though further experiments are required to test this hypothesis.

Numerous sequences coding for microRNA (miRNA) were also identified within the photosymbiotic bivalve NCRs. miRNAs play important roles in regulating gene expression by binding to targeted messenger RNAs (mRNAs), sometimes blocking translation or degrading mRNA (Du and Zamore 2007). The mitochondria are known to possess rich miRNA activities, although many of these miRNAs are imported from outside of the mitochondria and are not encoded in the mitogenome (Sripada *et al*. 2012). Having additional miRNAs directly encoded in the mitogenome may impact photosymbiotic bivalve gene expression, although much more in-depth research is needed to explore the roles of these miRNAs.

In addition to having regulatory elements, an extensive NCR could function as a hotbed for novel gene origination. Novel protein-coding genes could presently exist in those NCRs, as supported by the abundant ORFs identified within. Interestingly, many of the ORFs were overlapping with TEs, indicating that many of the TE fragments could still be transcribed. Overwhelming research have shown that TEs can be co-opted for host gene regulatory activities, such as adding regulatory elements (*e.g.*, promoter and enhancers), coding for transcriptional effector proteins, or providing gene silencing mechanisms (Fueyo *et al*. 2022). Given the numerous TE insertions in the photosymbiotic bivalves, it is highly possible that some of them have been co-opted for host mitochondrial functions.

Lastly, given that high frequency of discontinuous repeats might be selected for to prevent breaking of DNA secondary structures (Sebastian *et al*. 2021), some of the tandem repeats observed in this study are likely to form secondary structures, which are known to play important gene regulatory roles (Nardi *et al*. 2012). Although we observe highly divergent primary repeat sequences among different photosymbiotic bivalve lineages, it is possible that some of them form similar secondary structures, which can be conserved across taxa even though primary sequences are not (Sebastian *et al*. 2021).

## CONCLUSION

Our study sheds light on previously underappreciated yet ecologically significant aspects of animal mitogenome evolution: NCR structures and functional annotations. Our work using cardiids as examples implicates the potential associations between symbiotic lifestyles and mitogenome complexity. Apart from contributing to one of the largest mitogenomes amongst metazoans, the inflated major non-coding region in photosymbiotic bivalves, with either several independent origins and/or frequent rearrangements, have complex patterns in highly repetitive elements (nested tandem repeats) that require further characterizations. Many ORFs in the NCRs might contain more unrecognized genetic components but with hence far unknown functions, including active TE and other regulatory elements. While we have explored certain causes for having expanded NCRs, comparative analyses with active swimming or periodically oxygen-deprived pectinids (of which inflated NCRs have been frequently reported) within phylogenetics context would provide a more comprehensive understanding of how oxygen-related lifestyles may affect mitogenome evolution. With potentially important roles in ecological adaptation and mitogenome evolution, NCR and its structures in more animal species should be further explored.

## MATERIALS AND METHODS

### DNA Extraction and Genome Sequencing

Complete mitogenomes of six marine, photosymbiotic bivalve species (family Cardiidae) –– *Fragum fragum, F. whitleyi, F. sueziense* (subfamily Fraginae)*, Tridacna crocea, T. derasa,* and *T. gigas* (subfamily Tridacninae), were sequenced as part of the Aquatic Symbiosis Genomics Project by the Wellcome Sanger Institute. Detailed methods and genome statistics are described in the published genome notes (Table 1). In short, fresh tissue samples from living specimens were snap-frozen with liquid nitrogen and high molecular weight DNA was extracted. After shearing and purifying, PacBio HiFi circular consensus DNA sequencing libraries were constructed and sequenced on the PacBio SEQUEL II system (HiFi). The mitochondrial genome was assembled using MitoHiFi (Uliano-Silva *et al*. 2023). Mitogenomes of three non-photosymbiotic cardiids –– *Acanthocardia tuberculata* (subfamily Lymnocardiinae), *Cerastoderma edule* (subfamily Lymnocardiinae)*, Fulvia mutica* (subfamily Cardiinae) –– were obtained from NCBI Genbank (Table 1).

### NCR Verification through Illumina sequencing

Genomic DNA for two other distinct individuals of *F. fragum* and *F. whitelyi* collected from Guam (specimens deposited in the University of Colorado Museum of Natural History collection) were extracted using the E.Z.N.A.® Mollusc DNA Kit, according to the manufacturer’s instructions. Genomic libraries were prepared with the Nextera® XT DNA Library Prep Kit (Illumina®), and each sample was uniquely barcoded with Nextera® i5 and i7 dual index adapters. Library preparation and sequencing were performed at the University of Colorado BioFrontiers Institute Next-Generation Sequencing Facility. Samples were sequenced as paired-end 150 bp reads on the Illumina Novaseq S4 platform at Novogene. (Illumina raw reads from this study are available on NCBI GenBank BioProject PRJNA1162570). Reference-guided assembly mapping of Illumina short reads (150 bp) to PacBio mitogenome using *SAMtools* and *BCFtools* were then conducted to verify the presence, position and alignment of the major NCR (Danecek *et al*. 2021).

### Annotation and Comparative Analysis of Mitogenomes

Gene annotations for all species were performed and crosschecked using MITOS2 v2.1.8 on usegalaxy.org (Donath *et al*. 2019) and GeSeq on Chlorobox (Tillich *et al*. 2017). All protein-coding genes were also checked using NCBI ORFfinder to correctly identify the start and stop positions of their open reading frames. Transfer RNA genes were identified using tRNAscan-SE (Chan and Lowe 2019) and ARWEN (Laslett and Canbäck 2008); false positives were excluded after checking for their potential to form a typical tRNA clover-shaped secondary structure based on MFOLD (Zuker 2003). Comparative analysis of gene order and rearrangement were conducted via visual inspection and subsequently graphed in Adobe InDesign. Duplicated region and repeat patterns were identified and visualized using dotplots on NCBI Blast, and further detected using Tandem Repeat Finders v4.09 (Benson 1999). The Interspersed Repeat Protein Masking Based on Protein Similarity program on RepeatMasker is used to identify sequences that match transposable element (TE) encoded proteins (Smit *et al*. 2015). NCBI ORFfinder is further used to identify all ORFs within all mitogenomes (search parameters: minimal 75- nucleotide cut-off, invertebrate genetic code, “ATG” start codon and alternative initiation codons). Only ORFs that match TE/TE fragments, as well as a subset of major tandem repeats which falls within the following selection criteria are included in our main annotation dataset: 1. high period/consensus size, ≥100 nt OR 2. high copy number, ≥ 10, AND 3. High percent matches ≥ 98%. The main annotation dataset is then used for the linearized graphical representations of mitogenomes in this paper. Respective complete annotation lists are also made available as Data S1. Detailed graphical representations of the circular mitogenomes were visualized using OGDRAW (Greiner *et al*. 2019), and are individually included as Figs. S2a-S2i.

## ACKNOWLEGDMENT

Many thanks to the Aquatic Symbiosis Genomics Project for enabling large-scale long-read genomic sequencing of symbiotic life forms including our photosymbiotic cockles and clams. We would like to express our gratitude to Dr. Nolan C. Kane, Dr. Kyle Keepers and Emily Curcio for help and support in genomic sequencing and analyses, and Dr. Edward Chuong for insightful conversations pertaining to transposable elements. We are extending our appreciation to Kelly R. Martin, along with the University of Colorado Museum of Natural History, for the support in our use of specimens and databased records, and to Yu Kai Tan and Andressa Viol for aiding our polygon visualizations of mitogenome segments in R.

## AUTHOR CONTRIBUTIONS

Conceptualization: ADYT, JL Methodology: ADYT, RL, JL Investigation: ADYT Visualization: ADYT Supervision: JL Writing—original draft: ADYT, RL, JL Writing—review & editing: ADYT, RL, JL

## DATA AVAILABILITY

All data are available in the main text or the supplementary materials: Gene annotation datasets underlying this article are available in its online supplementary material; genomic sequences and Illumina raw reads used are publicly available on NCBI GenBank (see references in article and in Table 1). R codes for linearized visualization of mitogenomes will be shared on reasonable request to the corresponding author (these codes will be deposited in a Github repository before publication).

